# Ethanol Self-Administration Reduces mGlu2/3 Protein Expression in the Nucleus Accumbens and mGlu2/3 Activation Suppresses Binge Drinking in Male Mice

**DOI:** 10.64898/2026.03.18.712674

**Authors:** Cassandra G. Modrak, Sarah E. Holstein, April Kim, Elizabeth G. Shannon, Sara Faccidomo, Joyce Besheer, Clyde W. Hodge

## Abstract

**Background:** Alcohol use disorder (AUD) is associated with dysregulated glutamatergic signaling within mesocorticolimbic circuits that govern reinforcement and excessive ethanol intake. Group II metabotropic glutamate receptors (mGlu2/3) act primarily as presynaptic autoreceptors that regulate glutamate release. However, how voluntary alcohol intake alters mGlu2/3 expression within reward circuitry remains unclear.

**Methods and Results:** We examined mGlu2/3 protein expression following a history of voluntary ethanol exposure and operant self-administration and, in separate cohorts, assessed the functional impact of group II receptor modulation on binge-like ethanol intake. Male C57BL/6J mice received two weeks of home-cage access to sweetened ethanol or sucrose followed by 35 days of operant self-administration. Immediately after the final session, tissue punches from the nucleus accumbens (NAc), amygdala, and prefrontal cortex were collected for immunoblot analysis. mGlu2/3 expression was reduced in the NAc of ethanol exposed mice with no statistically significant changes detected in the amygdala or prefrontal cortex. In separate binge-drinking cohorts, systemic administration of the mGlu2/3 agonist LY379268 reduced binge-like ethanol intake in a limited-access home-cage drinking model, whereas positive allosteric modulation of mGlu2 receptors with LY487379 was ineffective.

**Conclusions:** Voluntary ethanol self-administration was associated with reduced mGlu2/3 protein expression in the NAc. Systemic pharmacological activation of group II receptors suppressed binge-like ethanol consumption. These findings identify mGlu2/3 dysregulation in a key reward-related brain region as a molecular adaptation associated with voluntary ethanol intake and demonstrate mechanistic regulation of binge drinking. This suggests that group II metabotropic glutamate signaling is a viable therapeutic target for treating AUD.

## Introduction

Alcohol use disorder (AUD) is characterized by dysregulation of excitatory neurotransmission within cortico-limbic circuits that govern reinforcement, motivation, and behavioral control (Koob, 2021, Koob and Volkow, 2016, Koob, 2003). Converging evidence from preclinical and clinical studies indicates that repeated ethanol exposure produces enduring glutamatergic neuroadaptations that extend beyond acute intoxication and withdrawal, contributing to excessive intake, loss of control, and relapse vulnerability (Gass and Olive, 2008). Among these circuits, the nucleus accumbens (NAc) functions as a critical integrative hub, processing convergent glutamatergic inputs to drive reward-related behavior and motivation (Kalivas, 2009, Kalivas et al., 2009, Besheer et al., 2010, Faccidomo et al., 2025c, Faccidomo et al., 2021).

Voluntary ethanol self-administration alters both presynaptic and postsynaptic components of glutamatergic signaling within the NAc (Faccidomo et al., 2025c, Woodward Hopf and Mangieri, 2018, Sidhpura et al., 2010, Kufahl et al., 2011). In particular, adaptations that elevate extracellular glutamate tone or enhance postsynaptic responsiveness are thought to promote ethanol reinforcement and escalation of intake (Faccidomo et al., 2025b, Zamudio et al., 2021, Cannady et al., 2017, Agoglia et al., 2015a, Griffin et al., 2014, Sidhpura et al., 2010). Growing evidence has identified impaired glutamate uptake via GLT-1 and altered cystine-glutamate exchange through system xc^-^ as key contributors to dysregulated glutamate homeostasis following ethanol and drug exposure (Reissner and Kalivas, 2010, Scofield et al., 2016, Griffin et al., 2021, Duclot et al., 2024). However, the specific molecular mechanisms that regulate active presynaptic glutamate release following voluntary, reinforcement-driven ethanol intake remain to be fully elucidated.

Group II metabotropic glutamate receptors (mGlu2 and mGlu3) are G_i/o_-coupled receptors that function as presynaptic autoreceptors and heteroreceptors to inhibit glutamate release (Schoepp, 2001, Testa et al., 1998). By constraining synaptic and extrasynaptic glutamate levels, mGlu2/3 receptors provide critical inhibitory control over excitatory transmission within reward-related circuits. Disruption of mGlu2/3 signaling has been linked to excessive glutamate release and maladaptive plasticity across models of synaptic transmission, neural plasticity, and substance use disorders (Lovinger and McCool, 1995, Gereau and Conn, 1995, Qian et al., 2019, Kalivas, 2009). Accordingly, pharmacological modulation of mGlu2/3 regulates ethanol intake, discrimination and relapse-like behavior (Vengeliene and Spanagel, 2022, Griffin et al., 2014, Cannady et al., 2011, Kufahl et al., 2011, Besheer et al., 2010, Sidhpura et al., 2010, Zhao et al., 2006, Tyler et al., 2024).

Despite strong pharmacological evidence implicating mGlu2/3 in ethanol-related behaviors, relatively little is known about how voluntary, non-dependent ethanol self-administration alters mGlu2/3 receptor expression in a brain region-specific manner. Evidence showing ethanol-induced alteration of mGlu2/3 expression relies primarily on non-contingent exposure (Meinhardt et al., 2013, Griffin et al., 2015), which addresses pharmacological impact but does not account for behavioral variables such as reinforcement history or the complex associative processes inherent in drug-seeking. Recent work further suggests that mGlu3 signaling within mesocorticolimbic circuits may contribute to ethanol reinforcement. Specifically, reduced mGlu3 protein expression in the NAc shell and prelimbic cortex has been identified in selectively bred alcohol-preferring (P) rats relative to control strains, indicating a genetically linked difference in mGlu3 signaling associated with elevated extracellular glutamate and enhanced ethanol reinforcement (Tan et al., 2026). Because voluntary ethanol self-administration engages learning, motivation, and presynaptic glutamatergic regulation, it may drive neuroadaptations distinct from those produced by passive exposure, which is a distinction that carries significant translational implications for modeling AUD.

Accordingly, the present study tested the hypothesis that voluntary ethanol self-administration alters mGlu2/3 receptor expression within specific reward-related brain regions. Group II metabotropic glutamate receptors (mGlu2 and mGlu3) are *Class C* G-protein coupled receptors that exist as dimers, with functional signaling requiring proper assembly of monomeric subunits into dimeric complexes (Pin and Bettler, 2016). Thus, alterations in monomeric availability or dimeric receptor abundance may reflect distinct but complementary forms of synaptic regulation, with implications for receptor signaling capacity, trafficking, and responsiveness to endogenous glutamate. To distinguish ethanol-associated molecular adaptations from nonspecific effects of operant learning or performance demands, data from ethanol self-administering mice were compared with sucrose-reinforced controls that showed no statistically significant reinforcer-group differences across multiple measures of operant performance. This design allowed molecular differences to be evaluated in the context of similar measured operant behavior while controlling for nonspecific features of training and reinforcement (Faccidomo et al., 2025c, Faccidomo et al., 2025b, Salling et al., 2016).

In a complementary functional approach, we examined whether systemic pharmacological modulation of group II metabotropic glutamate receptors alters binge-like ethanol intake using two mechanistically distinct compounds: the orthosteric mGlu2/3 agonist LY379268 and the mGlu2-selective positive allosteric modulator (PAM) LY487379. These experiments utilized a limited-access home-cage drinking procedure to evaluate receptor modulation under a pattern of binge-like ethanol consumption that has strong translational relevance (Thiele et al., 2014, Becker and Lopez, 2004). Comparison of LY379268 and LY487379 allowed evaluation of the behavioral effects of broad group II receptor activation relative to selective enhancement of mGlu2 signaling. Because the molecular and pharmacological studies were conducted in separate cohorts using different ethanol-exposure procedures, they provide complementary but independent information regarding mGlu2/3 expression and the effects of group II receptor modulation on ethanol-related behavior.

## Materials and methods

### Animals

Adult male C57BL/6J mice (n = 52) were obtained from The Jackson Laboratory (Bar Harbor, Maine) at 9 weeks of age upon arrival. Mice were group-housed for operant self-administration (n=16) and singly housed for binge drinking (n = 36) under standard laboratory conditions in clear polycarbonate cages (28 x 17 x 14 cm) with a stainless-steel wire top lid and corn cob bedding. This group size provides > 93% power to detect medium to large effect sizes for self-administration at α=0.05. Mice had *ad libitum* access to food and water except where noted. The vivarium was maintained at 21 ± 1^ο^C and 40 ± 2% humidity and a 12h:12h reverse light cycle (lights off at 0800). Mice acclimated to the vivarium for 1 week before initiation of experimental procedures. All procedures were conducted in accordance with the University of North Carolina-CH institutional and NIH guidelines.

### Operant Ethanol and Sucrose Self-Administration

Operant self-administration procedures were conducted as previously described (Faccidomo et al., 2025a, Hoffman et al., 2023, Hoffman et al., 2021, Faccidomo et al., 2021, Faccidomo et al., 2020, Faccidomo et al., 2015, Faccidomo et al., 2009). Operant conditioning chambers (Med Associates, St. Albans, VT, USA) were equipped with active and inactive levers and a recessed drinking cup. Presses on the active lever delivered a liquid reinforcer (0.014 ml/reinforcement) paired with a cue light and pump sound (800 ms). Presses on the inactive lever had no programmed consequence. Reinforcer deliveries were detected via photobeam activation in the drinking cup.

Separate groups of mice were given free home-cage access to water and either sweetened ethanol (9% ethanol + 2% sucrose; n = 8) or sucrose alone (2%; n = 8) for two weeks (24 h/day, 7 d/week) prior to operant training. Mice then completed four overnight (16 h) training sessions during which the response requirement increased from fixed-ratio 1 to fixed- ratio 4 (FR1–FR4), followed by 35 daily 1 h sessions under a fixed-ratio 4 (FR4) schedule with the same solutions serving as reinforcers. Previous work demonstrates that this model results in pharmacologically relevant blood ethanol levels after 1 h sessions and mice fully consume the reinforcing fluid without food or water restriction (Faccidomo et al., 2009) with comparable operant performance between ethanol- and sucrose-reinforcement conditions (Faccidomo et al., 2025c). Behavioral measures included responses on the active and inactive levers, percent active responding relative to total responses, number of reinforcers earned, and head entries into the drinking cup per reinforcer delivered.

### Tissue Collection and Immunoblotting

Tissue collection, processing and immunoblotting were conducted as previously described (Faccidomo et al., 2025c, Faccidomo et al., 2025b, Faccidomo et al., 2020, Faccidomo et al., 2018). Mice were sacrificed immediately following the 35th operant self-administration session. Tissue samples were collected using a 1.0-mm-diameter stainless-steel biopsy punch from 1.0-mm-thick coronal sections guided by the mouse brain atlas of Paxinos and Franklin (Paxinos and Franklin, 2001). Punches targeted the NAc (approximately AP +1.2 to +1.7 mm from bregma), amygdala (approximately AP −1.0 to −2.0 mm from bregma), and prefrontal cortex (PFC, approximately AP +1.7 to +2.7 mm from bregma), with punch placement centered over each region according to atlas-defined anatomical landmarks. Samples were then homogenized in an SDS buffer containing protease and phosphatase inhibitors (1X Halt Cocktail) and protein concentration was quantified using a BCA Assay (ThermoFisher). Protein samples (8 µg) were loaded into 4-15% Criterion TGX precast gels (Bio-Rad) and a standard gel electrophoresis protocol was run. Proteins were transferred to PVDF membranes using an iBlot^®^ Semi-Dry blotting system (Thermo Fisher Scientific).

Immunoblotting was conducted using antibodies against mGlu2/3 (rabbit anti-mGlu2/3, 1:1,000; Millipore Sigma) and GAPDH (mouse anti-GAPDH, 1:10,000; Advanced Immunochemical) as a loading control. The mGlu2/3 antibody was selected based on vendor validation for immunoblot detection of mGlu2/3, and our immunoblots produced bands consistent at the expected molecular weights of monomeric (∼106 kDa) and dimeric (∼200 kDa) mGlu2/3. Horseradish peroxidase-conjugated secondary antibodies (1:10,000; Jackson ImmunoResearch) and chemiluminescent detection reagents (Amersham ECL Prime; Cytiva) were used to visualize western blot signal. Immunoreactive bands were captured using an ImageQuant LAS4000 system (GE Healthcare Life Sciences).

Densitometric quantification was performed using ImageQuant TL software (GE Healthcare Life Sciences) by an investigator blinded to the experimental group with samples assigned to lanes in randomized order. Monomeric and dimeric mGlu2/3 bands were identified based on their expected molecular weights and quantified separately. Band densities were background-subtracted and normalized to the corresponding GAPDH loading-control signal from the same lane. Normalized values were then expressed relative to the mean of the sucrose control group within each brain region, with sucrose controls set to 100%. Monomeric and dimeric mGlu2/3 were quantified separately, and an overall mGlu2/3 composite index was calculated as the average of normalized monomeric and dimeric values for each sample [(M + D)/2]. Full immunoblots, including GAPDH loading-control bands for all analyzed samples, are provided in Supplemental Figure 1 to allow evaluation of band specificity, signal consistency, and quantification.

Individual samples were excluded for technical quality-control reasons including an inadequate GAPDH loading-control signal, insufficient tissue recovered during micropunch collection, or low protein yield after homogenization. Final sample sizes were NAc (Sucrose n = 8; EtOH n = 6), amygdala (Sucrose n = 6; EtOH n = 6), and PFC (Sucrose n = 7; EtOH n = 8). No samples were excluded on the basis of experimental outcome.

### Intermittent Binge Ethanol Drinking

After habituation to the vivarium, mice were acclimated to the limited-access binge-like drinking procedure by providing access to ethanol (20% v/v, n = 24) in place of water for 2 h per day for 2 consecutive days as previously reported (Gianessi et al., 2025, Agoglia et al., 2015b, Agoglia et al., 2015a, Holstein et al., 2011, Lowery et al., 2010). Following this acquisition period, mice were transitioned to the 4 h intermittent binge-access procedure, in which binge access drinking sessions, occurring 3 h into the dark cycle, were interspersed with alternating days of *ad libitum* access to water only. This procedure lasted 14 days, with ad libitum access to water occurring every other day. During ethanol access days, only ethanol was available for 4 h via a single 10-ml calibrated sipper tube. Using these tubes, fluid volume was measured to the nearest 0.1 ml at both 2- and 4 h time points without needing to remove the tube from the cage lid. For all drinking sessions, mice were weighed and returned to their home cages prior to fluid access. To account for potential leakage due to rack movement, sipper tubes were placed on empty cages during each session, and the average volume lost from these control tubes was subtracted from individual intake measurements. Ethanol intake was calculated as grams of ethanol per kilogram of body weight.

### Effects of mGlu2/3 Receptor Modulation on Binge-Like Ethanol Drinking

To assess the functional role of mGlu2/3 receptor modulation in binge-like ethanol consumption, male C57BL/6J mice (n = 12) received intraperitoneal injections of the selective mGlu2/3 agonist LY379268 (0, 0.3, 1.0, or 3.0 mg/kg) 30 min prior to ethanol access during the last four intermittent limited-access binge-drinking sessions. Drug testing was conducted on alternating drinking days (48 h between drug test sessions) using a within-subjects Latin square design to control for order effects. The 0 mg/kg condition served as the vehicle control, in which mice received an equivalent saline injection as part of the Latin square design. Dose selection and pretreatment interval were based on previous studies demonstrating the behavioral efficacy of LY379268 (Karlsson et al., 2008, Backstrom and Hyytia, 2005, Sidhpura et al., 2010). Ethanol intake (20% v/v) was measured 2 and 4 h after presentation of the drinking solution on each test day.

Sucrose intake was evaluated as a control to determine if the effects of LY379268 were specific to binge-like ethanol drinking or generalized to consumption of a non-ethanol palatable solution. A separate cohort of male C57BL/6J mice (n = 12) was trained using the same limited-access drinking procedure but received access to sucrose (1% w/v) instead of ethanol. The lowest effective dose of LY379268 identified in the ethanol study (0 or 1.0 mg/kg) was administered 30 min prior to sucrose access according to a counterbalanced within-subjects crossover design. Sucrose intake (mg/kg) was measured 2 and 4 h after presentation of the solution.

To compare orthosteric activation of mGlu2/3 receptors with selective positive allosteric modulation of mGlu2 receptors, a separate cohort of mice (n = 12) underwent the same binge-drinking procedures but received intraperitoneal injections of LY487379 (0, 3, 10, or 30 mg/kg), a selective mGlu2 positive allosteric modulator (PAM), 30 min before ethanol access. This experiment was conducted in parallel with the LY379268 study to determine whether selective enhancement of mGlu2 signaling was sufficient to alter binge-like ethanol intake. Drug testing occurred on alternating drinking days using a within-subjects Latin square design to control for order effects, with the 0 mg/kg dose serving as the vehicle condition. Ethanol intake was measured 2 and 4 h following presentation of the ethanol solution.

### Locomotor activity control

To determine whether pharmacologically induced reductions in ethanol intake could be attributed to nonspecific motor suppression, spontaneous locomotor activity was evaluated as previously reported (Faccidomo et al., 2015, Salling et al., 2008). The lowest effective dose of LY379268 (1.0 mg/kg) that reduced ethanol intake and the highest dose of the mGlu2-selective positive allosteric modulator LY487379 (30 mg/kg), which did not alter binge-like drinking, were tested in separate mice. The same mice used in respective binge-drinking experiments (n = 12 per compound, n = 6 per dose) were randomly assigned to vehicle or drug treatment (n = 6/treatment) for locomotor testing. One day following the last day of intermittent access (48 hours after the last 4-hr drinking session), mice received intraperitoneal injections of LY379268 (0 or 1.0 mg/kg) or LY487379 (0 or 30 mg/kg) 30 min prior to open-field testing. Locomotor behavior was assessed during one 2 h session in a Plexiglas open-field activity monitoring chamber (27.9 × 27.9 cm; Med Associates). Each chamber was equipped with two arrays of 16 pulse-modulated infrared photobeams that recorded horizontal (X-Y) ambulatory movements. The position of each mouse was sampled at 100-ms intervals, and distance traveled (cm) was calculated across the session in 10-min intervals using automated acquisition software.

### Drugs

LY379268 [(1R,4R,5S,6R)-4-amino-2-oxabicyclo[3.1.0]hexane-4,6-dicarboxylic acid] (Ascent Scientific, Princeton, NJ) was dissolved in 0.9% physiological saline. LY379268 is a highly selective group II mGlu receptor agonist with EC50 values of 2.69 nM for mGlu2 and 4.48 nM for mGlu3 subtypes, and shows no agonist or antagonist properties at mGlu1a, mGlu5a, or mGlu7 receptors when tested in a concentration range of (10,000 – 100,000 times the EC50 values for mGlu2/3 receptor activation (Monn et al., 1999).

LY487379 hydrochloride [2,2,2-Trifluoro-*N*-[4-(2-methoxyphenoxy)phenyl]-*N*-(3-pyridinylmethyl)ethanesulfonamide hydrochloride] (Tocris Bioscience, Minneapolis, MN) was suspended in carboxymethylcellulose (CMC 0.5% in 0.9% physiological saline). LY487379 is a subtype-selective mGlu2 PAM that potentiates glutamate-induced receptor signaling without intrinsic agonist activity. *In vitro* studies demonstrate that LY487379 enhances glutamate-stimulated [^35S]GTPγS binding with high potency at mGlu2 receptors (EC_₅₀_ ≈ 1.7 µM), while showing minimal activity at mGlu3 and other mGlu receptor subtypes, consistent with an allosteric mechanism dependent on endogenous glutamate (Johnson et al., 2003). All intraperitoneal (i.p.) injections were administered with a 27-gauge needle and an injection volume of 1ml/100g body weight.

Sucrose (1% w/v) was prepared in tap water and ethanol (20% v/v) was prepared by diluting 95% ethanol in tap water.

### Statistical Analysis

Behavioral and biochemical data were analyzed using parametric statistical tests implemented in GraphPad Prism. Statistical significance was set a priori at p < 0.05, and all tests were two-tailed. Immunoblot data were analyzed using independent-samples t-tests with Welch’s correction, which does not assume equal variance between groups. Variance comparisons are reported where relevant to aid interpretation of group differences, particularly for immunoblot measures showing unequal dispersion. Effect sizes and 95% confidence intervals are reported to facilitate interpretation of the magnitude and biological relevance of observed effects.

Operant self-administration data were analyzed using two-way RM-ANOVA, with reinforcer group as the between-subjects factor and session day as the repeated within-subjects factor. Comparison between ethanol- and sucrose-reinforced groups was evaluated prior to mGlu2/3 protein comparisons to ensure that molecular outcomes were interpreted in the context of comparable operant performance, including active lever responding, inactive lever responding, lever discrimination, and reinforcers earned. Drug effects on binge-like ethanol or sucrose intake were analyzed using repeated-measures ANOVA with dose and/or time as within-subjects factors, as appropriate for each experiment. Statistical outcomes were interpreted in conjunction with effect sizes and confidence intervals.

## Results

### Self-Administration: Comparison of Ethanol and Sucrose Groups

Separate groups of mice were trained to self-administer sweetened ethanol (n = 8) or sucrose (n = 8) during a 35-day baseline period (**Fig 1A**). The ethanol group consumed 0.793 ± 0.115 g/kg (mean ± SEM) per session during the final week. Ethanol- and sucrose-reinforced responding was comparable across groups, supporting the interpretation of effective behavioral matching prior to biochemical analyses. Beginning with the overnight sessions, there were no group differences between active (F(1, 14) = 0.15, p = 0.708) and inactive lever responses (F(1, 14) = 3.32, p = 0.090) (not shown). Operant performance across the 35-day baseline period is shown in **Fig 1B-G**. No significant main effects of reinforcer group were detected for active lever responses (F(1, 14) = 2.27, p = 0.158), inactive lever responses (F(1, 14) = 0.40, p = 0.536), percentage of responses on the active lever (F(1, 14) = 0.62, p = 0.443), reinforcers earned (F(1, 14) = 2.31, p = 0.151), total head entries (F(1, 14) = 3.39, p = 0.087), and head entries per reinforcer (F(1, 14) = 0.002, p = 0.790). A mixed-effects analysis found that both the EtOH (F(1, 14) = 23.31, p = 0.0003) and the sucrose-exposed mice (F(1, 14) = 12.96, p = 0.003) pressed significantly more on the active relative to the inactive lever, demonstrating lever discrimination in both groups. Lack of statistically significant group differences across these behavioral measures supports interpretation of subsequent molecular differences as associated with ethanol exposure rather than differences in operant performance. Tissue was prepared immediately after the last session for immunoblot analysis.

**Figure 1.**
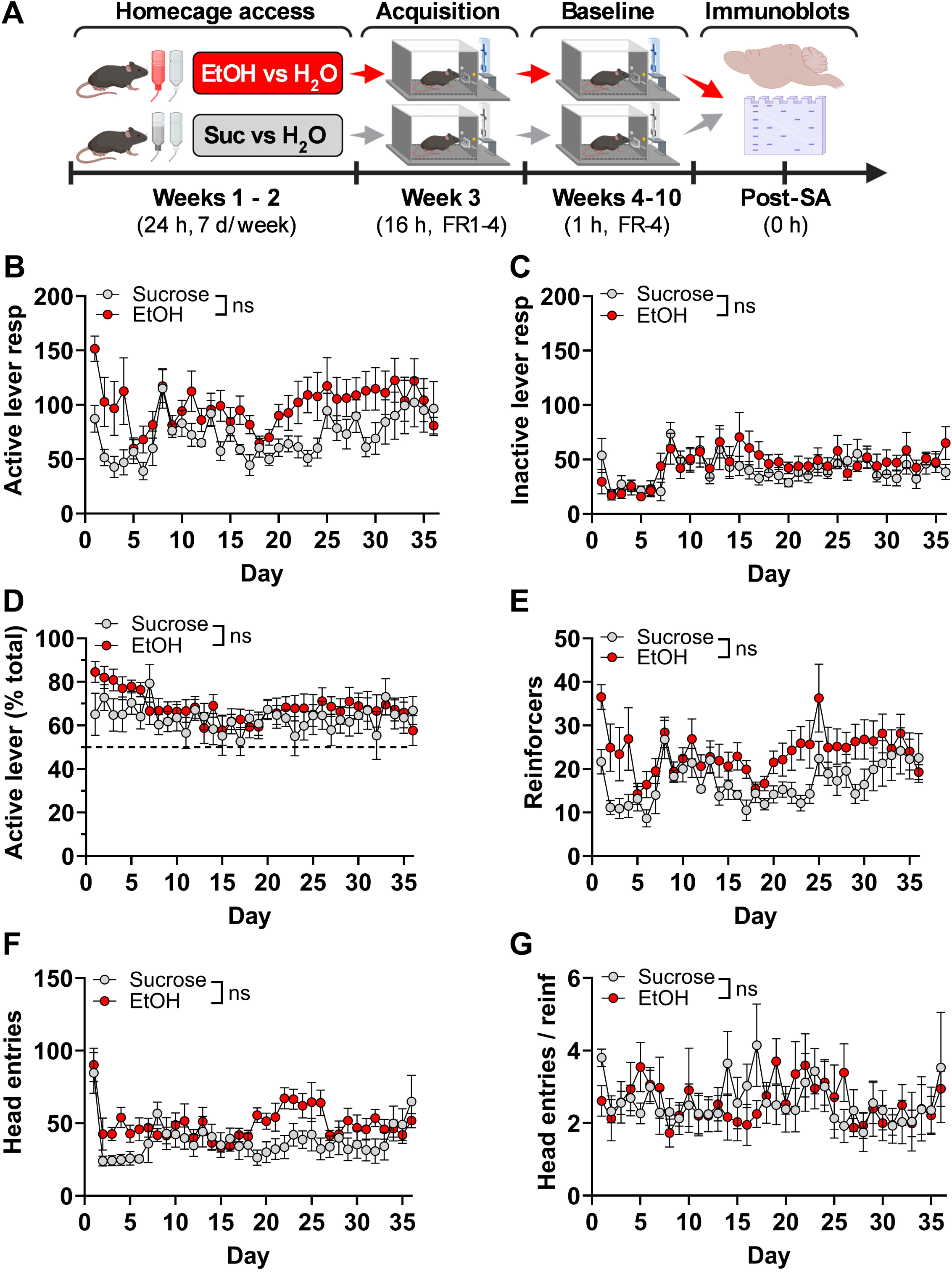
Parameters of operant ethanol and sucrose-only self-administration. **(A)** Experimental timeline for operant self-administration showing initial home-cage exposure (EtOH vs H_2_O two-bottle access), acquisition of operant behavior (EtOH or Sucrose - active vs inactive lever), and baseline for both reinforcement conditions culminating in tissue collection. **(B)** Total number of responses on the active lever for EtOH and Sucrose reinforcement plotted as a function of time (Day). (**C**) Total number of responses on the inactive lever. (**D**) Active lever responses as a percentage of total responses showing lever discrimination above 50% (*dashed line*). (**E**) Total number of reinforcers earned. (**F**) Total number of head entries into the delivery well. (**G**) Number of head entries per reinforcer earned as an index of consummatory behavior. Data are plotted as mean ± SEM for each day of the 35-day self-administration period; ***ns*** indicates no significant difference between EtOH and sucrose groups.

### Nucleus Accumbens: Downregulation of mGlu2/3 Protein Expression

#### mGlu2/3 Composite Index

To characterize overall mGlu2/3 protein expression in the NAc, monomeric and dimeric mGlu2/3 protein levels were combined into a composite index ((M + D) / 2), reflecting average expression across both molecular forms. Because monomeric and dimeric mGlu2/3 signals provide complementary measures of receptor protein expression, this composite index was used to assess the overall pattern of mGlu2/3 expression in addition to analyses of each molecular form separately. Sample sizes for NAc analysis were n = 8 (Suc) and n = 6 (EtOH). Two mice were excluded from the EtOH group due to insufficient protein yield (n = 1) and an incomplete tissue extraction (n = 1).

The mGlu2/3 composite index was significantly reduced in mice with a history of voluntary ethanol exposure and operant self-administration compared with sucrose controls (two-tailed independent-samples t test with Welch’s correction, t(9.52) = 2.25, p = 0.049, **Fig. 2A**). Mean composite mGlu2/3 expression was reduced by approximately 32% in the ethanol group relative to sucrose controls (EtOH: 67.7; Suc: 100.0). The magnitude of this effect was large (η² = 0.35), and the 95% confidence interval for the mean difference (−64.6 to −0.05) excluded zero. Variance did not differ significantly between groups (F(7,5) = 6.74, p = 0.052).

**Figure 2.**
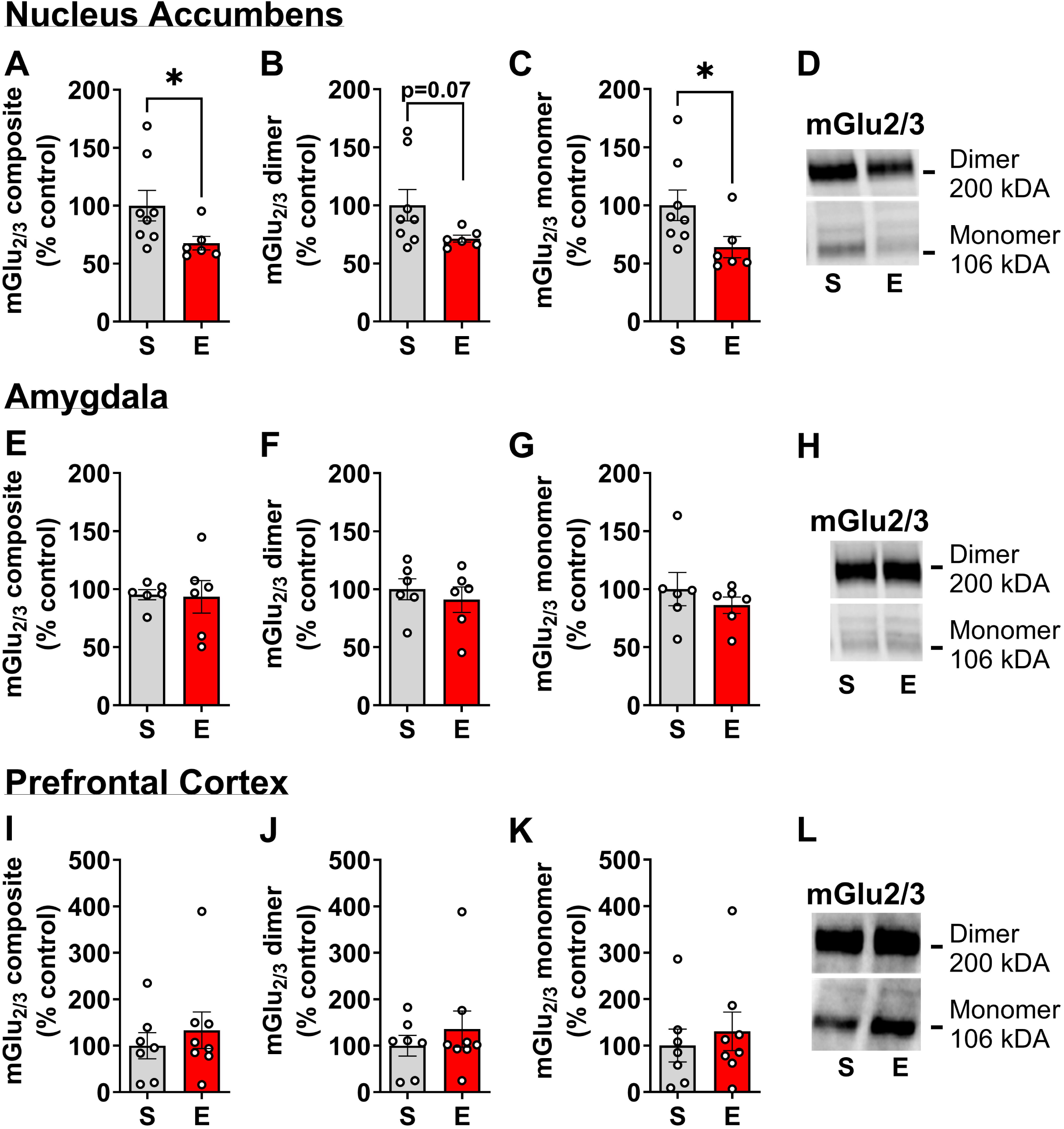
Operant ethanol self-administration reduces mGlu2/3 protein expression in the nucleus accumbens as compared to sucrose-only controls. (**A**) In the NAc, mGlu2/3 expression, calculated as the sum of normalized monomeric (M) and dimeric (D) mGlu2/3 protein levels ((M + D) / 2), was significantly reduced in ethanol self-administering mice compared with sucrose controls. (**B**) Dimeric mGlu2/3 protein expression in the NAc was reduced in the same direction but did not reach statistical significance. (**C**) Monomeric mGlu2/3 protein expression in the NAc was significantly reduced following operant ethanol self-administration. (**D**) Representative NAc immunoblots showing dimeric (∼200 kDa) and monomeric (∼106 kDa) mGlu2/3 bands and corresponding GAPDH loading controls. (**E – H**) Composite, dimeric, and monomeric mGlu2/3 protein expression and representative immunoblots from the amygdala, showing no ethanol-associated change. (**I – L**) Composite, dimeric, and monomeric mGlu2/3 protein expression and representative immunoblots from the PFC, showing no ethanol-associated change. Quantitative data were normalized to GAPDH, expressed relative to sucrose controls, and are shown as mean ± SEM. *p < 0.05 compared with sucrose controls, as determined by two-tailed Welch-corrected independent-samples t-tests. Sample sizes for each comparison are reported in the Results.

#### Dimeric mGlu2/3

Dimeric mGlu2/3 protein expression was reduced by approximately 29% in mice that self-administered ethanol relative to sucrose controls, although this effect did not reach statistical significance under two-tailed testing (EtOH: 71.3; Sucrose: 100.0; t(7.70) = 2.10, p = 0.07, **Fig 2B**). The magnitude of the effect was large (η² = 0.36), and the direction of change was concordant with the reductions observed for both monomeric mGlu2/3 expression and the total mGlu2/3 protein index. The 95% confidence interval for the mean difference included zero (-60.4 to 3.09), consistent with greater variability in the dimeric mGlu2/3 measure. Consistent with this interpretation, variance differed significantly between groups (F(7,5) = 26.2, p = 0.002), reflecting greater dispersion among sucrose-reinforced mice.

#### Monomeric mGlu2/3

Monomeric mGlu2/3 protein expression was approximately 36% lower in the ethanol-exposed group than in sucrose controls (EtOH: 64.0; Sucrose: 100.0; two-tailed Welch-corrected independent-samples t test, t(11.61) = 2.24, p = 0.045; Fig. 2C). The effect size was large (η² = 0.30), and variance did not differ significantly between groups (F(7,5) = 2.75, p = 0.28). Representative monomeric and dimeric mGlu2/3 immunoblots are shown in **Fig. 2D**, with full immunoblots provided in **Supplemental Fig. 1**.

### Amygdala: No Change in mGlu2/3 Protein Expression

No statistically significant differences were detected in mGlu2/3 protein expression between ethanol- and sucrose-exposed groups in the amygdala. The mGlu2/3 composite index did not differ between groups (two-tailed independent-samples *t* test with Welch’s correction, *t*(5.92) = 0.12, *p* = 0.91, **Fig 2E**; η² = 0.003; 95% confidence interval for the mean difference = -37.81 to 34.20). Similarly, no statistically significant group differences were detected for dimeric mGlu2/3 (t(9.65) = 0.64, p = 0.54, η² = 0.044; **Fig 2F**) or monomeric mGlu2/3 (t(7.32) = 0.87, p = 0.41, η² = 0.004; **Fig 2G**). Representative immunoblots are shown in **Fig 2H** and full immunoblots in Supplemental **Fig 1B**. Sample sizes were n = 6 (Suc) and n = 6 (EtOH), with two mice per group being excluded from analysis due to improper tissue extraction and one protein loading problem.

### Prefrontal Cortex: No Change in mGlu2/3 Protein Expression

No statistically significant differences between ethanol- and sucrose-exposed groups were detected in PFC mGlu2/3 protein expression. The mGlu2/3 composite index did not differ between groups (two-tailed Welch-corrected independent-samples t test, t(12.23) = 0.69, p = 0.50; **Fig. 2I**; η² = 0.037; 95% CI of the mean difference: −72.2 to 138.6). Similarly, no significant group differences were detected for dimeric mGlu2/3 (t(11.01) = 0.80, p = 0.44, η² = 0.055; **Fig. 2J**) or monomeric mGlu2/3 (t(12.92) = 0.56, p = 0.58, η² = 0.024; **Fig. 2K**). Representative immunoblots are shown in **Fig. 2L** and full immunoblots in **Supplemental Fig. 1**. Sample sizes were n = 7 (Sucrose) and n = 8 (EtOH), with one sucrose-exposed mouse excluded due to unsuccessful tissue extraction.

### Testing mGlu2/3 Regulation of Binge Ethanol Drinking

To determine whether pharmacological activation of mGlu2/3 receptors reduces binge-like ethanol consumption, the effects of systemic administration of the selective mGlu2/3 agonist LY379268 were evaluated during intermittent limited-access drinking sessions (**Fig 3A**). Ethanol intake was analyzed separately for the initial 2 h drinking period and for total intake across the full 4 h session.

**Figure 3.**
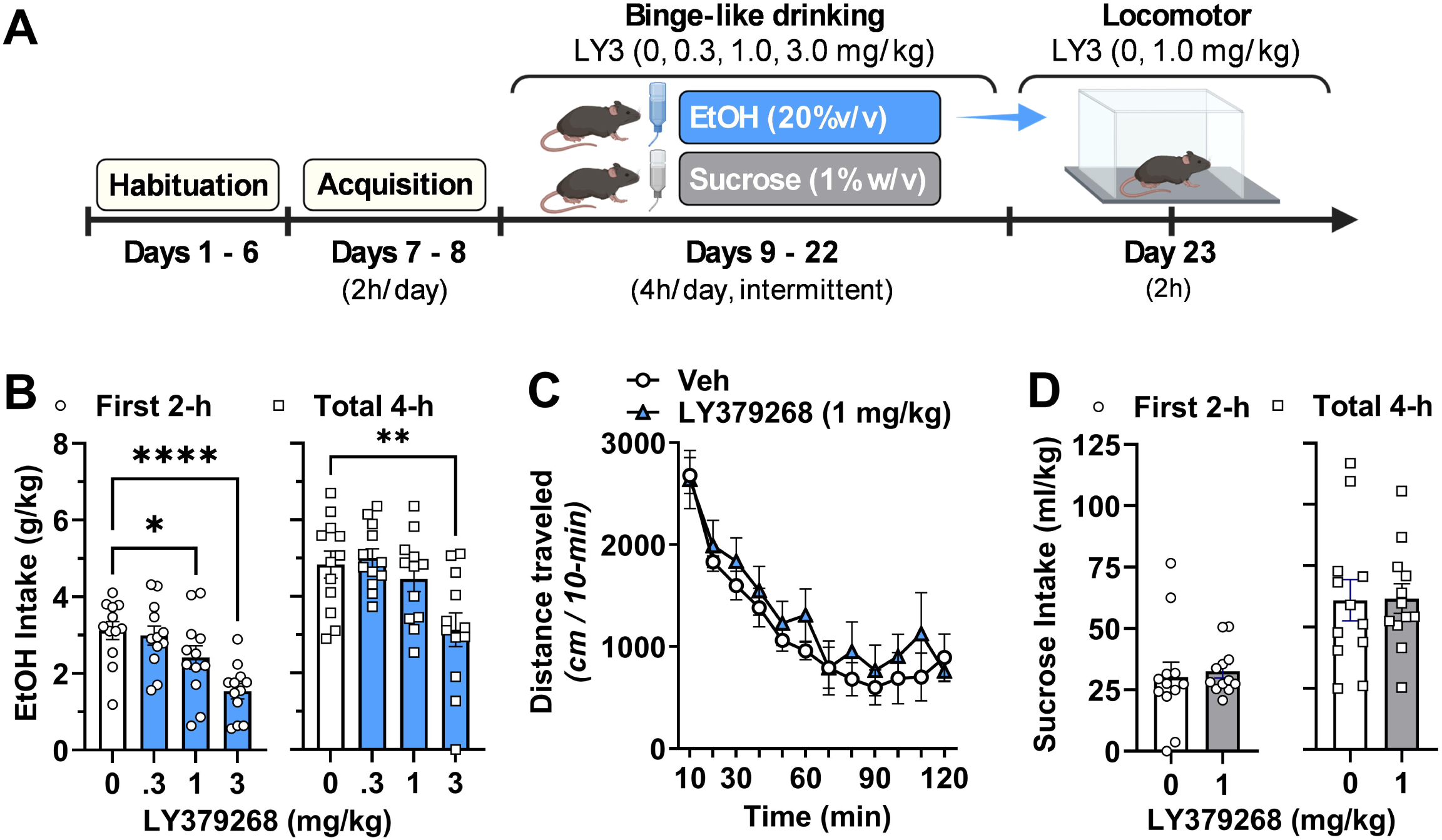
Pharmacological activation of mGlu2/3 receptors suppresses binge-like ethanol drinking. (**A**) Experimental timeline of binge drinking and locomotor procedures and LY379268 (LY3) administration. Both EtOH and Sucrose groups were initially habituated to the laboratory followed by initial acquisition of binge-like drinking (2h / day) and subsequent pharmacological testing of LY379268 (0 – 3.0 mg/kg) during 4h / day access periods. Finally, locomotor activity was tested in EtOH exposed mice after completion of drinking. Locomotor control testing was not conducted in sucrose mice due to the absence of effect of LY3 on sucrose intake. (**B**) Systemic administration of the mGlu2/3 agonist LY379268 dose-dependently reduced binge-like ethanol intake during the first 2 h of access (**left**) and total intake across the full 4 h session (**right**). Significant reductions relative to vehicle were observed at 1.0 and 3.0 mg/kg during the first 2-h, whereas only the highest dose reduced total 4 h intake. (**C**) In EtOH exposed mice, the lowest effective dose of LY379268 (1.0 mg/kg) did not alter spontaneous locomotor activity in an open-field test. Distance traveled declined across time bins, reflecting normal habituation, with no main effect of drug treatment or Time x Treatment interaction. (**D**) LY379268 (1.0 mg/kg) did not affect binge-like sucrose intake during either the first 2 h (**left**) or the full 4 h session (**right**), indicating that reductions in ethanol intake were not due to nonspecific suppression of consummatory behavior. All data are expressed as mean ± SEM. Symbols indicate significant differences relative to vehicle as determined by repeated-measures ANOVA with Holm-Šídák post hoc comparisons (*p < 0.05, **p < 0.01, ****p < 0.0001).

#### Ethanol Intake: First 2 Hours

Systemic administration of LY379268 produced a significant reduction in ethanol intake during the first 2 h of access (repeated-measures one-way ANOVA, *F*(3,33) = 9.22, *p* = 0.0001; **Fig 3B, left**). The magnitude of this effect was large (R² = 0.46), indicating that LY379268 dose accounted for approximately 46% of the variance in early-session ethanol intake. Post hoc Holm-Šídák comparisons showed that both the 1.0 and 3 mg/kg doses significantly reduced ethanol intake relative to vehicle (*p* = 0.044 and *p* < 0.0001, respectively).

#### Ethanol Intake: Total 4 Hours

A significant effect of LY379268 dose was also observed on total ethanol intake measured across the full 4 h drinking session (repeated-measures one-way ANOVA, *F*(3,33) = 6.80, *p* = 0.0011; **Fig 3B, right;** R² = 0.38). Post hoc analyses indicated that the highest dose of LY379268 (3.0 mg/kg) significantly reduced total ethanol intake relative to vehicle (*p* = 0.0016), whereas the 1.0 mg/kg dose did not significantly alter total intake across the 4 h session (*p* = 0.425).

Further analysis of ethanol consumed specifically during the 2–4 h interval revealed no significant effect of LY379268 (repeated-measures one-way ANOVA with Greenhouse–Geisser correction, F(2.27, 24.91) = 0.60, p = 0.578). These data indicate that systemic activation of mGlu2/3 receptors suppressed binge-like ethanol intake, with the strongest effects observed during the initial 2 h phase of drinking.

#### Locomotor Activity: Lack of Nonspecific Effects of LY379268

To determine whether the reductions in ethanol intake produced by LY379268 during the first 2 h of the binge drinking session could be attributed to nonspecific locomotor effects, spontaneous locomotor activity was assessed following systemic administration of the lowest effective dose of LY379268 (1.0 mg/kg) that reduced ethanol binge-like drinking. Distance traveled was measured across a 2 h open-field session and analyzed as a function of time and drug treatment. Locomotor activity showed a robust main effect of time, reflecting normal exploration and habituation across the test session (two-way repeated-measures ANOVA, *F*(11,110) = 27.02, *p* < 0.0001; **Fig 3C**). In contrast, there was no main effect of LY379268 treatment (*F*(1,10) = 0.55, *p* = 0.477), nor was there a significant time x treatment interaction (*F*(11,110) = 0.48, *p* = 0.911). The 95% confidence interval for the mean difference between vehicle- and LY379268-treated mice (-662.2 to 332.6) spanned zero. These findings indicate that the lowest effective dose of LY379268 did not alter spontaneous locomotor activity or habituation under these test conditions.

#### Sucrose Intake: Lack of Effect of LY379268

To determine whether the reduction in ethanol intake produced by LY379268 reflected a nonspecific suppression of consummatory behavior, the lowest effective dose that reduced ethanol intake (1.0 mg/kg) was tested on binge-like sucrose consumption. Systemic administration of LY379268 (1.0 mg/kg) did not significantly alter sucrose intake during the first 2 h of access compared with vehicle treatment (paired *t* test, *t*(11) = 0.52, *p* = 0.614; **Fig 3D, left**; partial η² = 0.024; 95% CI: -7.83 to 12.66. Similarly, LY379268 did not significantly affect total sucrose intake measured across the full 4 h session (paired *t* test, *t*(11) = 0.11, *p* = 0.914; **Fig 3D, right;** partial η² = 0.001; 95% CI: -13.06 to 14.44). This suggests that the suppressive effects of LY379268 (1 mg/kg) on ethanol intake are not attributable to generalized reductions in consummatory behavior.

### Testing mGlu2-Selective Regulation of Binge Ethanol Drinking

To determine whether selective potentiation of mGlu2 receptors is sufficient to reduce binge-like ethanol intake, the effects of systemic administration of the mGlu2-selective positive allosteric modulator LY487379 were evaluated across a range of doses (0, 3, 10, and 30 mg/kg) on binge-like ethanol intake followed by a locomotor activity control test (**Fig 4A**). In contrast to the suppressive effects observed with the mGlu2/3 agonist LY379268, LY487379 did not significantly alter ethanol intake during either the initial 2 h drinking interval or the total 4 h session.

**Figure 4.**
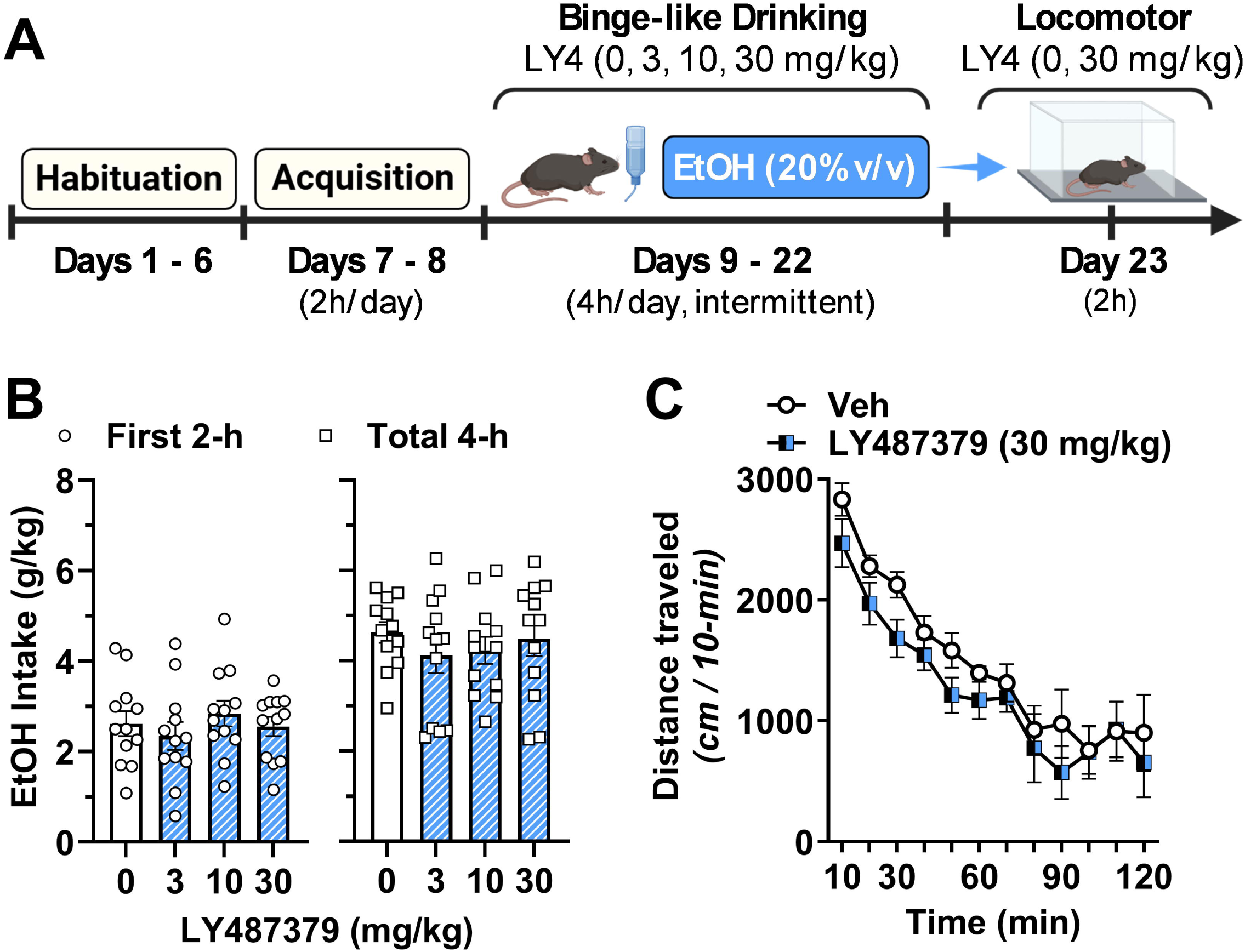
Positive allosteric modulator of mGlu2 receptors is ineffective at suppressing binge-like ethanol drinking. (**A**) Experimental timeline showing habituation, initial acquisition of EtOH binge drinking, pharmacological testing of LY487379, and locomotor control testing of the highest (ineffective) dosage of LY487379. Sucrose controls were not tested due to the lack of effect on EtOH binge drinking or locomotor activity. (**B**) Systemic administration of the mGlu2-selective PAM LY487379 (0-30 mg/kg) did not alter ethanol intake during the first 2 h (**left**) or across the full 4 h session (**right**), indicating that selective potentiation of mGlu2 receptors was insufficient to suppress binge-like drinking. (**C**) The highest dose of LY487379 (30 mg/kg) did not affect spontaneous locomotor activity, with normal habituation observed across the 2 h open-field session and no main effect of treatment. All data are expressed as mean ± SEM.

#### Ethanol Intake: First 2 Hours

During the first 2 h of binge-like ethanol drinking, repeated-measures ANOVA found no main effect of LY487379 treatment on ethanol intake (F(3,33) = 0.56, p = 0.65; **Fig 4B, left**; R² = 0.048), and post hoc Holm-Šídák comparisons indicated no differences between vehicle and any LY487379 dose (all adjusted p values > 0.86).

#### Ethanol Intake: Total 4 Hours

Similarly, analysis of total 4 h ethanol intake showed no significant effect of LY487379 treatment (F(3,33) = 0.98, p = 0.41; **Fig 4B, right;** R² = 0.082), and no pairwise comparisons between vehicle and drug doses reached significance (all adjusted p values ≥ 0.35). Across both time points, ethanol intake values remained comparable across all LY487379 doses, indicating that selective mGlu2 potentiation did not influence binge-like ethanol consumption.

##### Locomotor Activity: Lack of Nonspecific Effects of LY487379

To assess whether the highest LY487379 dose produced nonspecific behavioral suppression, locomotor activity was evaluated following administration of LY487379 (0 or 30 mg/kg). Two-way repeated-measures ANOVA revealed a robust main effect of time, reflecting normal habituation across the 2 h session F(11,110) = 33.94, p < 0.0001; **Fig 4C**), but no main effect of LY487379 treatment (F(1,10) = 1.61, p = 0.23) and no time x treatment interaction (F(11,110) = 0.50, p = 0.90). These findings indicate that LY487379 did not alter spontaneous locomotion at the highest dose tested on binge-like drinking.

## Discussion

The results of this study show that voluntary self-administration of sweetened ethanol was associated with reduced mGlu2/3 protein expression in the NAc as compared to sucrose-only controls. This effect was observed following a 35-day baseline of operant self-administration during which no statistically significant group differences were detected across multiple behavioral measures of ethanol- and sucrose-reinforced responding, which reduces the likelihood that the molecular findings reflected systematic differences in learning, memory or operant performance. No changes in mGlu2/3 expression were detected in the amygdala or PFC, suggesting anatomical specificity of this ethanol-induced adaptation within reward-related circuitry.

Given the established role of mGlu2/3 receptors as presynaptic autoreceptors that regulate glutamate release (Schoepp, 2001), reduced receptor expression in the NAc may have implications for inhibitory regulation of excitatory transmission and glutamate homeostasis. Immunoblot analysis of NAc tissue revealed significant reductions in the mGlu2/3 composite index and monomeric mGlu2/3 expression in the ethanol group relative to sucrose controls, together with a trend toward reduced dimeric expression. Because group II metabotropic glutamate receptors function as dimers (Pin and Bettler, 2016), changes across these molecular forms may reflect alterations in the available receptor protein pool and receptor organization. However, the present immunoblot measurements do not directly establish receptor signaling capacity or the functional state or subtype composition of the detected receptor complexes. T

The ethanol-induced reduction in mGlu2/3 expression observed in the present study is consistent with prior findings showing that ethanol exposure and excessive intake are associated with a hyperglutamatergic state within the NAc. Chronic intermittent ethanol vapor exposure (Griffin et al., 2014) and intermittent voluntary home-cage intake (Pati et al., 2016) both increase basal extracellular glutamate levels in the NAc across rodent models. This effect persists beyond acute withdrawal and is thought to promote excessive drinking and relapse vulnerability. Pharmacological studies further suggest that reducing glutamatergic tone within the NAc, either by enhancing glutamate uptake or by activating presynaptic mGlu2/3 autoreceptors, attenuates ethanol intake, particularly in dependent or high-drinking animals (Griffin et al., 2014). Moreover, systemic or site-specific activation of mGlu2/3 receptors using the selective mGlu2/3 agonist LY379268 reliably suppresses ethanol self-administration, cue-induced reinstatement, stress-induced relapse, and ethanol-related discriminative stimulus effects across a range of rat models (Backstrom and Hyytia, 2005, Zhao et al., 2006, Cannady et al., 2011, Jaramillo et al., 2015). Because the NAc serves as a major integrative hub for reward-related glutamatergic signaling (Scofield et al., 2016), mGlu2/3 expression within this region may be particularly sensitive to voluntary ethanol consumption. Alternatively, mGlu2/3 adaptations within amygdala and PFC may emerge under conditions of dependence, abstinence, or relapse, or may follow a different temporal profile than that examined in the present study.

Complementary behavioral experiments showed that systemic administration of LY379268 reduced binge-like ethanol intake, indicating that pharmacological activation of mGlu2/3 receptors is sufficient to suppress ethanol consumption under these conditions. Although ethanol-induced reductions in mGlu2/3 protein expression were observed in the NAc but not amygdala or PFC, systemic LY379268 would be expected to activate group II receptors throughout the brain. Thus, the observed regional pattern of mGlu2/3 reduction should not be interpreted as indicating that the behavioral effects of LY379268 are mediated exclusively within the NAc. Rather, activation of intact mGlu2/3 receptors across corticolimbic circuitry may contribute to the suppression of ethanol drinking, with reduced mGlu2/3 expression in the NAc identifying one neuroadaptation associated with voluntary ethanol self-administration.

An important finding from this experiment is that LY379268 (1.0 mg/kg) reduced ethanol intake during the initial 2 h of access without significantly altering sucrose consumption or spontaneous locomotor activity. Previous studies have reported that LY379268 can reduce sucrose seeking, cue-induced responding, or motivation under operant or reinstatement procedures, and in some cases suppress locomotor activity at higher doses or following site-specific administration (Bossert et al., 2006, Uejima et al., 2007, Windisch and Czachowski, 2018, Grimm et al., 2023). However, Bossert et al. (2006) also found no effect of LY379268 on sucrose self-administration, whereas studies reporting sucrose-related effects employed different behavioral procedures, species, or higher drug doses. Additionally, intra-accumbens or intra-amygdala administration of LY379268 has been associated with nonspecific motor suppression depending on dose and region (Cannady et al., 2011, Besheer et al., 2010). However, the present control experiments support behavioral specificity of the 1.0 mg/kg effect on early-session ethanol intake. Behavioral specificity was not directly assessed at 3.0 mg/kg, which reduced both early-session and cumulative 4-h ethanol intake. Analysis of ethanol consumed specifically during the 2-4 h interval revealed no significant effect of LY379268, indicating that effects were selective to the initial 2-h phase of drinking.

In contrast to the efficacy of the mGlu2/3 agonist LY379268, the mGlu2-selective PAM LY487379 did not alter binge-like ethanol intake. This pharmacological dissociation is of interest because positive allosteric modulators enhance receptor responses to endogenous glutamate rather than directly activating the receptor and therefore depend on endogenous transmitter tone and receptor availability (Conn et al., 2009). Thus, reduced mGlu2/3 expression in the NAc following alcohol intake may have contributed to the lack of efficacy observed with LY487379. Conversely, LY379268 remained effective in suppressing binge-like ethanol intake, consistent with prior demonstrations of agonist efficacy across models of excessive ethanol consumption and dependence (Zhao et al., 2006, Griffin et al., 2014). Previous work has also shown that systemic LY379268 administration but not the selective mGlu2 PAM BINA reduced ethanol seeking in rats (Windisch and Czachowski, 2018), providing further insight into pharmacological distinctions between these compounds as they relate to ethanol consumption. A parallel sucrose control experiment with LY487379 would have provided a useful assessment of whether mGlu2-selective PAM efficacy depends upon neuroadaptations associated with ethanol exposure. However, because LY487379 did not alter ethanol intake, this specificity control was not subsequently evaluated. Thus, interpretation of the LY487379 findings is limited to the conclusion that mGlu2-selective positive allosteric modulation was ineffective under the conditions tested.

Several limitations should be considered when interpreting these results. First, the molecular experiment included two weeks of voluntary home-cage ethanol exposure before operant training, and tissue was collected immediately following the final operant session. The observed NAc reduction therefore cannot be attributed uniquely to the operant self-administration phase and may reflect pretraining ethanol exposure, cumulative voluntary ethanol intake, ethanol consumed during the final session, or interactions among these factors. Because tissue was collected immediately following the final operant session, it also remains possible that acute ethanol exposure contributed to the observed reduction in mGlu2/3 expression. Future studies incorporating longitudinal sampling across abstinence and relapse-related time points will be important for determining the stability and behavioral relevance of this adaptation and for distinguishing acute from persistent neuroadaptations. Second, although reduced mGlu2/3 expression may have implications for glutamate signaling, direct functional measures of glutamate transmission were not assessed. Integrating molecular analyses with *in vivo*microdialysis or electrophysiological approaches will be necessary to establish a causal link between reduced mGlu2/3 availability and altered glutamate dynamics following voluntary ethanol intake. Third, although pharmacological activation of mGlu2/3 receptors was sufficient to suppress binge-like ethanol drinking, the present experiments did not determine whether endogenous mGlu2/3 signaling is required for regulating ethanol consumption. Future studies employing receptor antagonists, negative allosteric modulators, or genetic approaches will be necessary to address this question. Finally, the present study focused exclusively on male mice. Although prior work demonstrates robust sex differences in ethanol intake and motivation with conserved glutamatergic regulation of ethanol reinforcement across sexes, including uniform sensitivity to AMPAR antagonism (Faccidomo et al., 2025a), it remains unknown whether ethanol-induced downregulation of NAc mGlu2/3 expression is similarly conserved in females. Addressing this question will be critical for determining whether mGlu2/3-related neuroadaptations to low-dose operant ethanol self-administration represent a shared molecular substrate or exhibit sex-specific features, which may be relevant to precision therapeutic strategies.

In conclusion, the present findings add to a growing literature implicating group II metabotropic glutamate receptors in ethanol-related behaviors and identify reduced mGlu2/3 expression within the NAc as a neuroadaptation associated with voluntary ethanol self-administration. From a translational perspective, these findings highlight group II metabotropic glutamate receptors as a signaling system that may be relevant to the development and maintenance of excessive binge-like ethanol intake. Although the receptor subtype-specific mechanisms remain to be determined, the present results demonstrate that pharmacological activation of group II mGlu receptors is sufficient to suppress binge-like ethanol drinking. Together, these findings support continued investigation of mGlu2/3-targeted interventions as potential therapeutic approaches for alcohol use disorder.

## Data availability

The data that support the findings of this study are available from the corresponding author upon reasonable request.

