## Supplementary figures and images for "Ethanol Self-Administration Reduces mGlu2/3 Protein Expression in the Nucleus Accumbens and mGlu2/3 Activation Suppresses Binge Drinking in Male Mice" (from another version of this preprint: version 2, 2026-07-10)

### Supplemental Figure 1

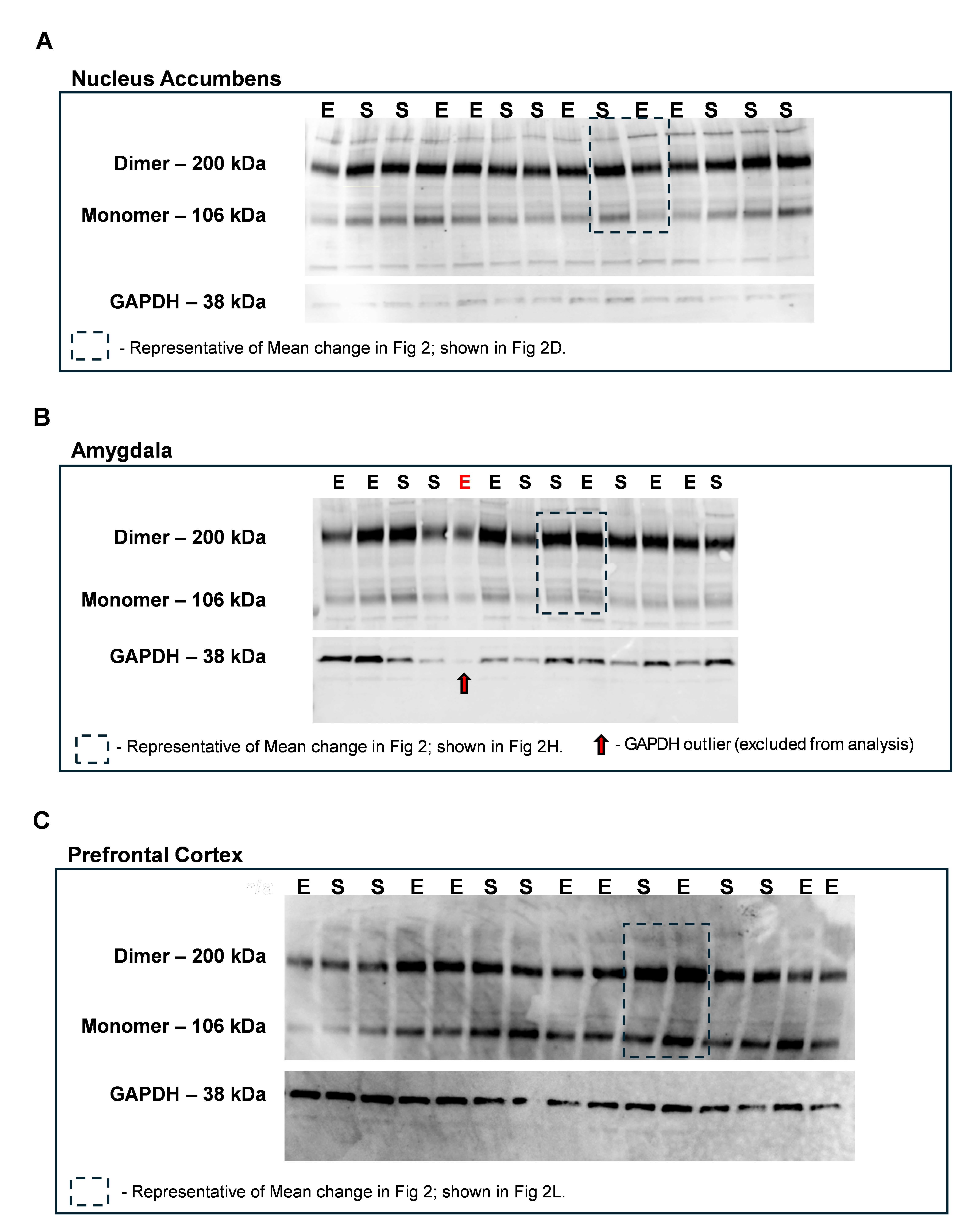
